# Settlement, growth and survival of eastern oysters on alternative reef substrates

**DOI:** 10.1101/010793

**Authors:** Seth J. Theuerkauf, Russell P. Burke, Romuald N. Lipcius

## Abstract

Restoration of the native eastern oyster (*Crassostrea virginica*) has been severely hindered by the dwindling supply and rising costs of fossil and new oyster shell for use in reef restoration. Consequently, emphasis has shifted to the use of alternative oyster reef materials, which need to be tested for their effectiveness as settlement substrate. Furthermore, low recruitment of wild larvae has also impeded restoration, indicating a need to assess the potential of field setting of cultured larvae. We experimentally examined oyster settlement, growth and survival on unconsolidated oyster shell, oyster shell embedded vertically in concrete, and concrete Oyster Castles® in field and mesocosm experiments. In addition, we examined settlement success of cultured larvae in the mesocosm experiment. In the field experiment, juvenile recruitment was 3x higher on castles and unconsolidated shell than on embedded shell. Castles retained 4x the number of oysters and hosted 5x the biomass than embedded shell, and retained 1.5x the oysters and hosted 3x the biomass than unconsolidated shell. The proportion of live oyster recruits on castles was 1.5x that on both embedded and unconsolidated shell. In the mesocosm experiment (90-d post-larval deployment), the castles recruited, retained, and hosted an oyster biomass 4x higher than that of unconsolidated and embedded shell. This study confirms that artificial reef materials such as Oyster Castles® are suitable alternative substrates for oyster restoration, and remote setting of larvae onto conditioned substrate can be effective under controlled environmental conditions. Future restoration efforts should consider use of alternative reef substrates and field setting of larvae to maximize oyster recruitment, while simultaneously minimizing the cost of reef restoration.

## 1. Introduction

Chesapeake Bay’s native oyster, *Crassostrea virginica* (Gmelin 1791), is an ecosystem engineer that performs critical ecological functions, including water filtration, sediment stabilization, and provision of nursery habitat for juveniles of diverse fish and invertebrate species (Kennedy et al., 1996). Prior to European colonization of North America, the native oyster population of Chesapeake Bay was described as being so abundant that “they lay as thick as stones” throughout the Bay and its tributaries. As a result of overfishing, disease, and poor water quality, the native oyster population of Chesapeake Bay currently stands at less than 1% of its historic population size (Rothschild et al., 1994; Wilberg et al., 2011). Additionally, human activities on land have increased the flow of sediment into the estuaries, which have weakened physiological health, lowered fecundity and raised mortality of oysters (Newell, 1988; Rothschild et al., 1994; Lenihan et al., 1999). Exacerbating the situation, the physical profile of reefs has been leveled by fishers exploiting oyster reefs (Rothschild et al., 1994), which places oysters lower in the water column where water flow is reduced and sediment accumulation rates are highest, thereby suffocating oysters (Newell, 1988; Lenihan et al., 1999).

Efforts to restore native oyster populations have been extensive but largely ineffectual or unresolved (Ruesink et al., 2005; Kennedy et al., 2011). However, recent successful restoration efforts with natural shell reefs and alternative materials have defined promising approaches (Lipcius and Burke, 2006; Taylor and Bushek, 2008; Powers et al., 2009; Schulte et al., 2009; La Peyre et al., 2014). In Chesapeake Bay as in other locations, availability and cost of reef substrate remains a significant hindrance to restoration progress (Kennedy et al., 2011). For instance, construction of sanctuary reefs in the Great Wicomico River oyster reef network required extensive use of dredged shell derived from buried fossil shell deposits (Schulte et al., 2009) at an estimated cost of nearly US$10,000 per ha per cm of reef. Cheaper substrates such as crushed concrete, limestone, and porcelain toilets have been used as alternative oyster reef materials (Soniat et al., 1991; Haywood III et al., 1999), but until recently their effectiveness has not been adequately tested against oyster shell (La Peyre et al., 2014). In addition, enhancement of oyster recruitment through larval setting remotely on such structures has not been investigated experimentally in the eastern oyster.

Alternative substrate may be preferentially settled upon by oyster larvae due to the large amount of surface area of a suitable chemical composition available for larval settlement. Additionally, larval settlement may be facilitated by the ability of these alternative substrates to mimic the three-dimensional structure of natural oyster reefs. The three-dimensional nature of many alternative substrates allows the oysters to be oriented above the benthic floor and to escape some of the sedimentation and predation faced by those that settle on thin layers of oyster shell.

In 2008, the Allied Concrete Corporation, in conjunction with The Nature Conservancy, developed an alternative reef substrate–Oyster Castle®. This prefabricated substrate features a parapet shape at the top of each block and is composed of limestone gravel, concrete, and crushed oyster shell, all of which have been used in various forms to construct artificial oyster reefs. Hence, Oyster Castle® reefs can serve as a model system to examine the utility of alternative substrates in oyster reef construction. Moreover, due to the stackable nature of the block and low cost of ingredients, three-dimensional reef structures of variable heights can be constructed easily and cost-effectively.

The objective of this study was to test the efficacy of alternative oyster reef substrates and remote setting of larvae in field and mesocosm experiments. Specifically, we compared settlement, size, and survival of oysters on Oyster Castle®, unconsolidated oyster shell, and oyster shell embedded vertically in concrete as a function of natural recruitment and artificially enhanced recruitment through remote setting.

## 2. Materials and Methods

### 2.1. Experimental reef substrates

Three reef substrates were tested in mesocosms and in the intertidal zone of the York River along the Virginia Institute of Marine Science (VIMS) in Gloucester Point, Virginia. As part of a balanced analysis of variance (ANOVA) randomized block design (Underwood, 1997), three sets of experimental blocks (larval subsidy) and three sets of control blocks (wild recruitment) were created using (i) four Oyster Castles® (hereafter referred to as OC; Figure 1a), (ii) one tray of loose, unconsolidated oyster shell (hereafter referred to as OS; Figure 1b) containing 16 shells (~75 mm shell height) per 0.25 m x 0.25 m quadrant, and (iii) one tray of vertically-embedded oyster shell (hereafter referred to as ES; Figure 1c) in a 0.5 m x 0.5 m base of concrete (Quikrete™ underlayment concrete) per block. ES served as a control for substrate vertical relief, and all substrates occupied a bottom area of 0.5 m x 0.5 m. Substrates were conditioned in the intertidal zone for two months to ensure that the physicochemical and biological (development of a biofilm) characteristics of the substrate surface were suitable for larval settlement (Bonar et al., 1990).

**Figure 1:**
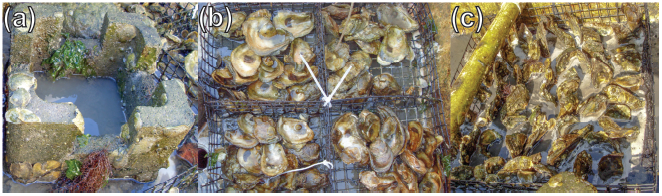
Photographs of the experimental reef substrates utilized in the mesocosm and field experiments, including (a) Oyster Castle®, (b) unconsolidated oyster shell, and (c) oyster shell embedded vertically in concrete.

### 2.2. Larval deployment

Hatchery-reared eyed oyster larvae were obtained from the VIMS Aquaculture Genetics and Breeding Technology Center in Gloucester Point, Virginia. Approximately six million oyster larvae were concentrated into a larval ball and stored overnight in cheesecloth at *~*5 °C. Immediately prior to release, the larvae were removed from cold storage, allowed to equilibrate to ambient air temperature (~20 min), and then immersed in a beaker containing a fixed volume (2 L) of river water. Light stirring was employed to evenly distribute larvae in solution and then divided across both the field and mesocosm experimental blocks (~950,000 larvae per block).

### 2.3. Mesocosm experiment

After the conditioning period, the experimental and control blocks were enclosed within three mesocosm tanks (Figure 2). An air supply system was customized to ensure that the mesocosms remained well-oxygenated and that ambient, flow-through river water was circulated throughout each mesocosm. Prior to larval introduction, each mesocosm was partitioned with silt fence to separate experimental blocks from control blocks. Following a standard aquaculture industry protocol, a single coat of petroleum jelly was liberally applied to deter settlement of larvae on the mesocosm surface (Congrove et al., 2008). Prior to larval release, water flow in the mesocosms was halted and airflow was reduced.

**Figure 2:**
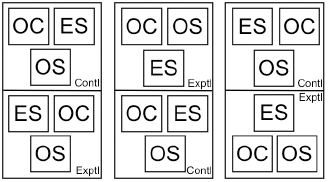
Layout of experimental and control blocks within the mesocosm experiment. Lines between blocks within mesocosms indicate silt fence barriers. ES = embedded oyster shell, OS = oyster shell, OC = Oyster Castle®.

### 2.4. Field experiment

The reef substrates were placed in the intertidal zone along the northern shore of the York River for conditioning and the duration of the experiment (Figure 3). After conditioning, the experimental blocks were enclosed with silt fence for a one week larval settlement period, and the control blocks remained exposed.

**Figure 3:**
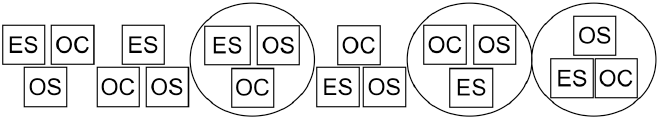
Layout of experimental and control blocks within the field experiment. Circles indicate silt fence enclosures surrounding experimental blocks. ES = embedded oyster shell, OS = oyster shell, OC = Oyster Castle®.

### 2.5. Sampling

At fixed sampling intervals (15, 45, and 90 d), oyster settlement, shell height, and survival were recorded. At each sampling interval, samples were randomly selected (Microsoft Excel™ Random Number Generator) from one of the four quadrants within each replicate. Shell height was measured with calipers and mortality determined by gaping oyster spat or presence of oyster scars on the substrate.

### 2.6. Statistical analysis

Analysis of variance (ANOVA) models were employed for both the field and mesocosm experiments using the 90-d sampling data. Ninety days was selected as the appropriate sampling interval for statistical analyses because this duration allowed for settlement of larvae, post-settlement mortality, and juvenile growth (Osman and Abbe, 1994; Burke, 2010). Response variables included (i) Total Recruits [total number of live and dead recruits], (ii) Live Recruits, (iii) Live Recruit Biomass [ash-free dry oyster tissue mass], and (iv) Proportion of Live Recruits [live recruits divided by total recruits]. The two fixed factors were Reef with three levels [OC, OS, ES] and Larval Subsidy with two levels [wild recruitment only, wild recruitment + larval subsidy]. Location was a blocking factor with three levels. Levene’s test was used to test the assumption of homogeneity of variance. The Student–Newman–Keuls (SNK) test was used to assess significant differences between factor levels.

Live oyster biomass, as ash-free dry mass (AFDM), was computed using the following power function (Lipcius unpubl data):

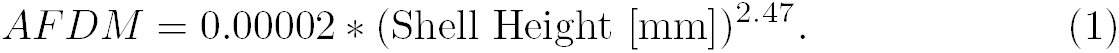

Akaike’s Information Criterion (AIC) within an Information Theoretic approach (Anderson, 2008) was used to evaluate five models (Table 1). *AIC*_*c*_ (a second-order bias correction estimator), Δ_*i*_ (a weighted measure of each model relative to the best fitting model), and *ω_i_* (model probability) were used to compare model (*g*_*i*_) fits. ANOVA tests were used to assess the goodness of fit of competing models.

**Table 1:** Parameters for the candidate linear regression models (response variables = total recruits, live recruits, live recruit biomass, and proportion of live recruits). *k* = number of parameters, including variance (*σ*^2^) as a parameter.

| Model | Description | $k$ | Effects | | | |
| --- | --- | --- | --- | --- | --- | --- |
|  |  |  | Subsidy | Reef | Subsidy x Reef | Block |
| $g_1$ | Full | 8 | x | x | x | x |
| $g_2$ | Interaction | 7 | x | x | x | |
| $g_3$ | Additive | 5 | x | x | | |
| $g_4$ | Reef | 4 | x | | | |
| $g_5$ | Subsidy | 3 | | x | | |

The effect of recruit size on survival was analyzed with a log-linear model using the frequencies of live and dead recruits as the response variable. Factors included recruit size (> or < 30 mm shell height), larval subsidy (yes, no), reef type (ES, OS, OC), and experiment (field, mesocosm). There were significant interactions including experiment as a factor, so the log-linear models were conducted separately for the field and mesocosm experiments. Effects were integrated into the models with stepwise addition of effects using AIC as the criterion. All analyses were run using the R statistical software package (R Core Team, 2014).

## 3. Results

### 3.1. Mesocosm experiment

Model *g*_*3*_, the additive model including Reef and Subsidy (Table 1), provided the best fit for Total Recruits (AIC *ω_i_* = 0.93), Live Recruits (AIC *ω_i_* = 0.93), and Recruit Biomass (*ω_i_* = 0.90); Table S1 provides model selection results for all models from the mesocosm experiment. For these three variables, the Reef and Subsidy factors were always significant (Table 2). In addition, neither the global model (*g*_*1*_) nor the interaction model (*g*_*2*_) improved the fit significantly better than model *g*_*3*_ (*F* test, *P* > 0.27 for all three variables).

**Table 2:** Analysis of variance results for model *g*_*3*_, the additive model with Reef and Subsidy as factors, from the mesocosm experiment. Although model *g*_*4*_ was the best fitting model for Proportion of Live Recruits, we present the results for model *g*_*3*_ because it shows the non-significant effect of Subsidy, and the results for models *g*_*3*_ and *g*_*4*_ did not differ in the significance of the factors.

| Treatment | MS | df | $F$ | $P$ |
| --- | --- | --- | --- | --- |
| *(a) Total recruits* |  |  |  |  |
| *Subsidy* | 2838 | 1 | 10.75 | 0.005 |
| *Reef* | 11757 | 2 | 44.53 | < 0.0005 |
| *Error* | 264 | 14 |  |  |
| *(b) Live recruits* |  |  |  |  |
| *Subsidy* | 2763 | 1 | 10.63 | 0.006 |
| *Reef* | 11045 | 2 | 42.52 | < 0.0005 |
| *Error* | 260 | 14 |  |  |
| *(c) Live recruit biomass* |  |  |  |  |
| *Subsidy* | 187.5 | 1 | 9.87 | 0.007 |
| *Reef* | 162.0 | 2 | 8.53 | 0.003 |
| *Error* | 19.0 | 14 |  |  |
| *(d) Proportion of live recruits* |  |  |  |  |
| *Subsidy* | 0.00011 | 1 | 0.85 | 0.37 |
| *Reef* | 0.0017 | 2 | 13.28 | 0.0006 |
| *Error* | 0.00013 | 14 |  |  |

The magnitude and direction of the Reef and Subsidy effects were equivalent for Total Recruits and Live Recruits (Table 3). To be concise, we only portray the patterns for Live Recruits (Figure 4). Live recruit density (Figure 4a) and biomass (Figure 4c) were substantially and significantly (SNK test, p < 0.05) higher on OC than on OS; those on OS were significantly higher than those on ES (SNK test, p < 0.05). Recruit density and biomass were more than twice as high on OC than on OS; those on OS were nearly twice as high as on ES (Figure 4a,c, Table 3). Recruit density was about 50% higher in the larval subsidy treatments (Figure 4b, Table 3), while biomass was more than twice as high (Figure 4d, Table 3). These differences were due to the earlier settlement and growth of cultured juveniles over wild juveniles, which resulted in a greater fraction of juveniles greater than 30 mm shell length with larval subsidy (Figure 5).

**Table 3:** Parameter estimates for model *g*_*3*_, the additive model with Reef and Subsidy as factors, from the mesocosm experiment. The value for the Intercept represents the mean for the No Subsidy/Embedded Shell treatment. Other values represent the additional effect sizes due to the specific treatments, and are additive.

| Parameter | Estimate | SE | $t$ | $P$ |
| --- | --- | --- | --- | --- |
| *(a) Total recruits* |  |  |  |  |
| *Intercept* | 14.94 | 7.66 | 1.95 | 0.07 |
| *Subsidy* | 25.11 | 7.66 | 3.28 | 0.005 |
| *Reef: Castle* | 84.33 | 9.38 | 8.99 | < 0.0005 |
| *Reef: Shell* | 18.83 | 9.38 | 2.01 | 0.06 |
| *(b) Live recruits* |  |  |  |  |
| *Intercept* | 15.11 | 7.60 | 1.99 | 0.07 |
| *Subsidy* | 24.78 | 7.60 | 3.26 | 0.006 |
| *Reef: Castle* | 81.50 | 9.31 | 8.76 | < 0.0005 |
| *Reef: Shell* | 17.50 | 9.31 | 1.88 | 0.08 |
| *(c) Live recruit biomass* |  |  |  |  |
| *Intercept* | 0.41 | 2.05 | 0.20 | 0.84 |
| *Subsidy* | 6.45 | 2.05 | 3.14 | 0.007 |
| *Reef: Castle* | 9.94 | 2.52 | 3.95 | 0.001 |
| *Reef: Shell* | 2.35 | 2.52 | 0.93 | 0.37 |
| *(d) Proportion of live recruits* |  |  |  |  |
| *Intercept* | 0.998 | 0.005 | 185.90 | < 0.0005 |
| *Subsidy* | 0.004 | 0.005 | 0.92 | 0.37 |
| *Reef: Castle* | -0.026 | 0.007 | -4.01 | 0.001 |
| *Reef: Shell* | -0.032 | 0.007 | -4.81 | 0.0003 |

**Figure 4:**
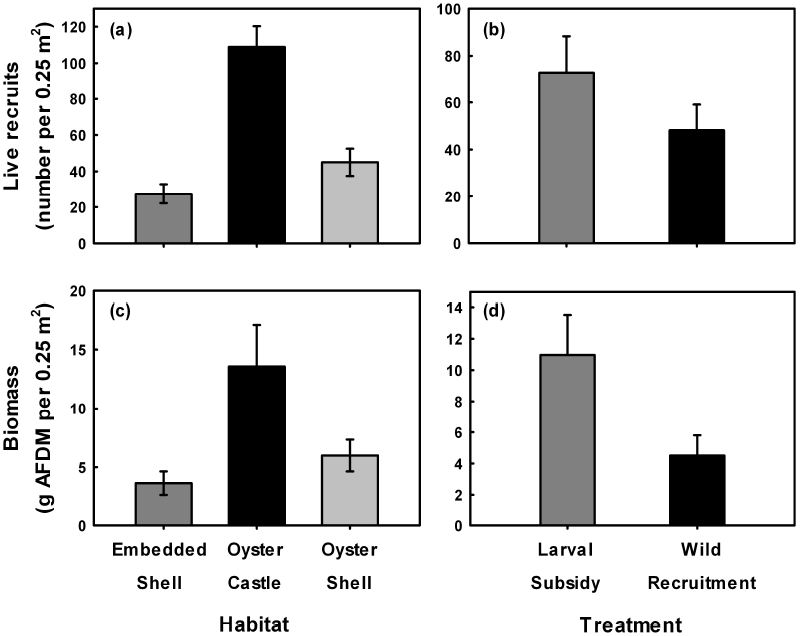
(a) Mean live recruits and (c) mean live recruit biomass on the various substrates, and (b) mean live recruits and (d) mean live recruit biomass in the experimental and control blocks, in the mesocosm experiment. Vertical bars represent one standard error of the mean.

**Figure 5:**
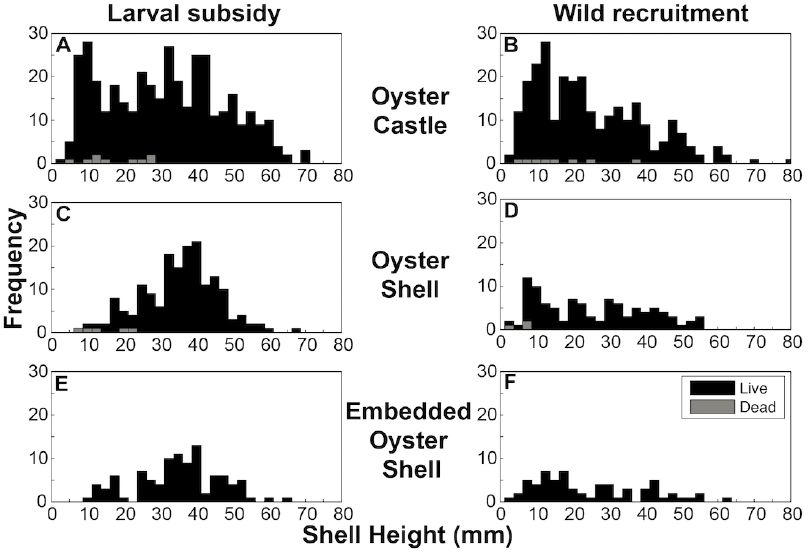
Size frequency histogram of live and dead oysters on Oyster Castles® (a & b), oyster shell (c & d), and embedded shell (e & f) in the mesocosm experiment.

For the Proportion of Live Recruits, model *g*_*4*_ –the model that only included Reef (Table 1)–provided the best fit (AIC *ω_i_* = 0.81), although model *g*_*3*_ also had substantial support (AIC *ω_i_* = 0.19); Table S1 provides model selection results for all models from the mesocosm experiment. The Reef factor was highly significant, whereas the effect of Subsidy was not significant (Table 2). In addition, neither the global model (*g*_*1*_) nor the interaction model (*g*_*2*_) improved the fit significantly better than model *g*_*4*_ (*F* test, *P >* 0.53).

There was very high survival in all treatments (Figure 5). The proportion of live recruits was nearly 0.998 in the ES treatment, and only dropped to 0.972 and 0.966 in the OC and OS treatments, respectively (Table 3). Although the effect of Reef was statistically significant (Table 2), the effect sizes were very small (Table 4) and not likely to be biologically significant (Figure 5).

**Table 4:**
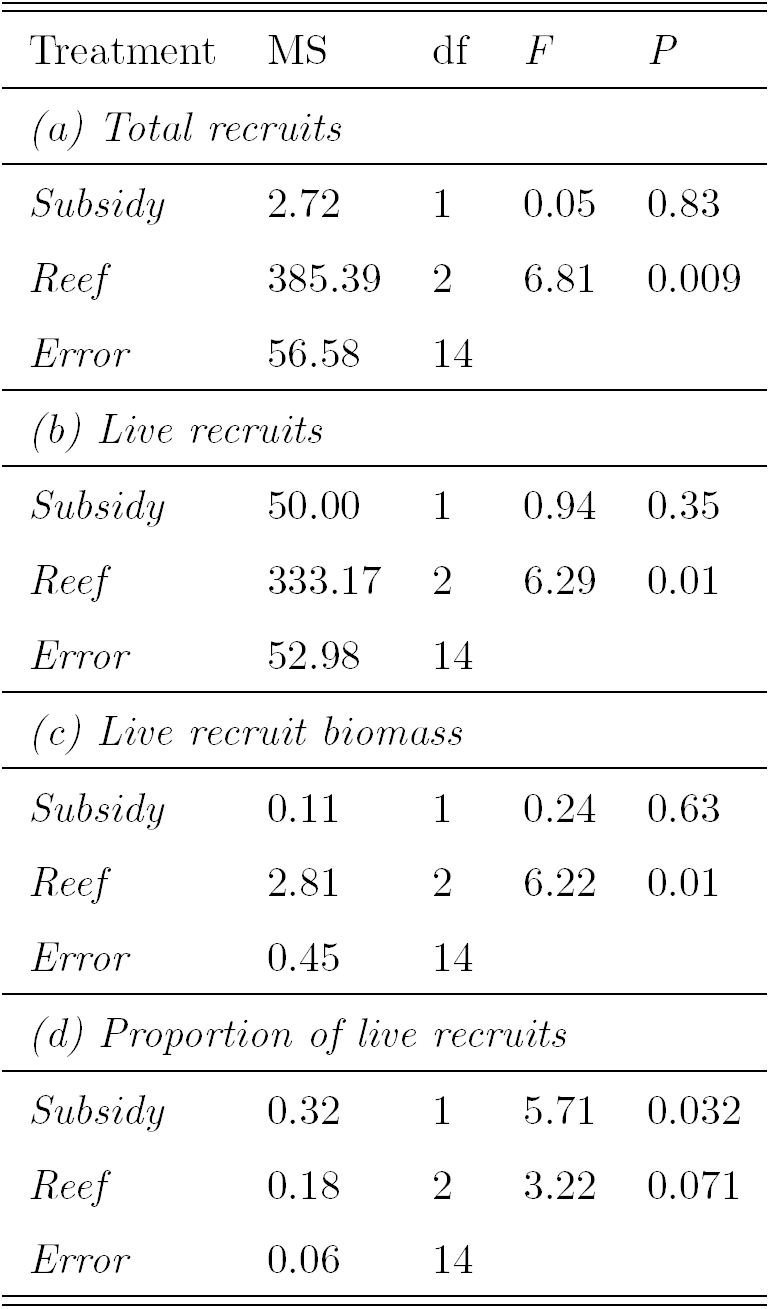
Analysis of variance results for model *g*_*3*_, the additive model with Reef and Subsidy as factors, from the field experiment. Although model *g*_*4*_ was the best fitting model for Total Recruits, Live Recruits, and Live Recruit Biomass, we present the results for model *g*_*3*_ for consistency.

| Treatment | MS | df | $F$ | $P$ |
| --- | --- | --- | --- | --- |
| *(a) Total recruits* |  |  |  |  |
| *Subsidy* | 2.72 | 1 | 0.05 | 0.83 |
| *Reef* | 385.39 | 2 | 6.81 | 0.009 |
| *Error* | 56.58 | 14 |  |  |
| *(b) Live recruits* |  |  |  |  |
| *Subsidy* | 50.00 | 1 | 0.94 | 0.35 |
| *Reef* | 333.17 | 2 | 6.29 | 0.01 |
| *Error* | 52.98 | 14 |  |  |
| *(c) Live recruit biomass* |  |  |  |  |
| *Subsidy* | 0.11 | 1 | 0.24 | 0.63 |
| *Reef* | 2.81 | 2 | 6.22 | 0.01 |
| *Error* | 0.45 | 14 |  |  |
| *(d) Proportion of live recruits* |  |  |  |  |
| *Subsidy* | 0.32 | 1 | 5.71 | 0.032 |
| *Reef* | 0.18 | 2 | 3.22 | 0.071 |
| *Error* | 0.06 | 14 |  |  |

### 3.2. Field experiment

We experienced storm conditions during our attempt to release hatchery-reared larvae in the field, which precluded a strong effect of larval subsidy in the field experiment (Table 4). Model *g*_*4*_, the model that only included Reef (Table 1), provided the best fit for Total Recruits (AIC *ω_i_* = 0.86), Live Recruits (AIC *ω_i_* = 0.78), and Recruit Biomass (*ω_i_* = 0.84); Table S2 provides model selection results for all models from the field experiment. For these three variables, the Reef factor was always significant (Table 4). In addition, neither the global model (*g*_*1*_) nor the interaction model (*g*_*2*_) improved the fit significantly better than model *g*_*3*_ (*F* test, *P >* 0.34 for all three variables).

Total recruit density (Figure 6a) was approximately equal (SNK test, p >> 0.05) on OC and OS; these were significantly higher than those on ES (SNK test, p < 0.05). Total recruit density was about threefold higher on OC and OS than on ES (Table 5a). Live recruit density (Figure 6b) and biomass (Figure 6c) were significantly higher on OC than on OS, and those on OS were significantly higher than those on ES (SNK test, p < 0.05). Live recruit density and biomass were about fivefold higher on OC than on ES; on OS they were slightly greater than those on ES (Table 5b,c).

**Table 5:** Parameter estimates for model *g*_*3*_, the additive model with Reef and Subsidy as factors, from the field experiment. The value for the Intercept represents the mean for the No Subsidy/Embedded Shell treatment. Other values represent the additional effect sizes due to the specific treatments, and are additive.

| Parameter | Estimate | SE | $t$ | $P$ |
| --- | --- | --- | --- | --- |
| *(a) Total recruits* |  |  |  |  |
| *Intercept* | 7.11 | 3.55 | 2.01 | 0.06 |
| *Subsidy* | 0.78 | 3.55 | 0.22 | 0.83 |
| *Reef: Castle* | 14.50 | 4.34 | 3.34 | 0.005 |
| *Reef: Shell* | 13.17 | 4.34 | 3.03 | 0.009 |
| *(b) Live recruits* |  |  |  |  |
| *Intercept* | 3.50 | 3.43 | 1.02 | 0.33 |
| *Subsidy* | 3.33 | 3.43 | 0.97 | 0.35 |
| *Reef: Castle* | 14.83 | 4.20 | 3.53 | 0.003 |
| *Reef: Shell* | 8.67 | 4.20 | 2.06 | 0.06 |
| *(c) Live recruit biomass* |  |  |  |  |
| *Intercept* | 0.26 | 0.32 | 0.82 | 0.43 |
| *Subsidy* | 0.15 | 0.32 | 0.49 | 0.63 |
| *Reef: Castle* | 1.26 | 0.39 | 3.24 | 0.006 |
| *Reef: Shell* | 0.16 | 0.39 | 0.41 | 0.69 |
| *(d) Proportion of live recruits* |  |  |  |  |
| *Intercept* | 0.49 | 0.11 | 4.42 | 0.0006 |
| *Subsidy* | 0.27 | 0.11 | 2.39 | 0.03 |
| *Reef: Castle* | 0.29 | 0.14 | 2.12 | 0.05 |
| *Reef: Shell* | -0.02 | 0.14 | -0.15 | 0.88 |

**Figure 6:**
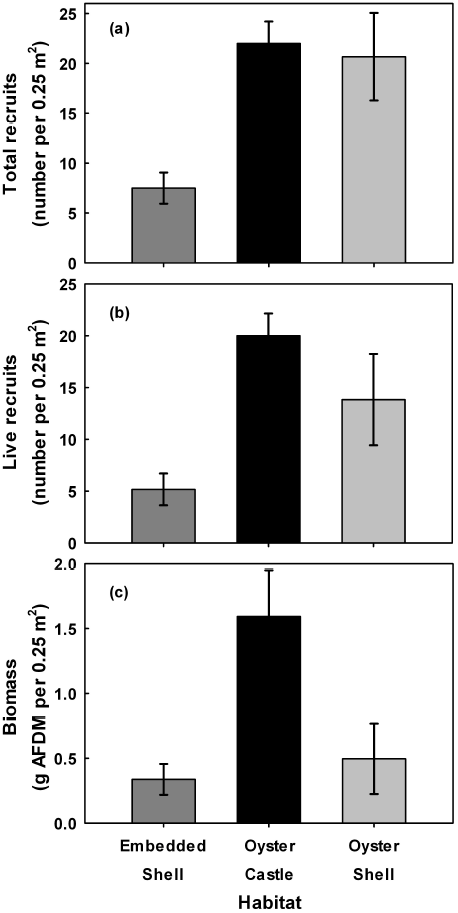
(a) Mean total recruits, (b) mean live recruits, and (c) mean live recruit biomass on the various substrates in the field experiment. Vertical bars represent one standard error of the mean.

For the Proportion of Live Recruits, models *g*_*3*_ (AIC *ω_i_* = 0.38) and *g*_*5*_ (AIC *ω_i_* = 0.49) provided reasonable fits to the data. We selected model *g*_*3*_ due to its higher explanatory power as indicated by the *r*^2^ values and due to the statistically significant effects of Reef and Subsidy (Tables 4, 5d). In addition, neither the global model (*g*_*1*_) nor the interaction model (*g*_*2*_) improved the fit significantly better than model *g*_*3*_ (*F* test, *P* > 0.47 for all three variables); Table S2 provides model selection results for all models from the field experiment.

There was moderate to high survival of recruits in all treatments (Figure 7). The proportion of live recruits was significantly higher with larval subsidy (Table 4), which increased survival by 47% (Figure 7a, Table 5d). Survival was also significantly higher on OC than on OS and ES (SNK test, p < 0.05), ranging from about 60% on OS and ES to 91% on OC (Figure 7b, Table 5d). Mortality occurred primarily in juveniles less than 30 mm shell length (Figure 8), but it did not produce a greater fraction of juveniles greater than 30 mm shell length, unlike the pattern for the mesocosm experiment (Figure 5).

**Figure 7:**
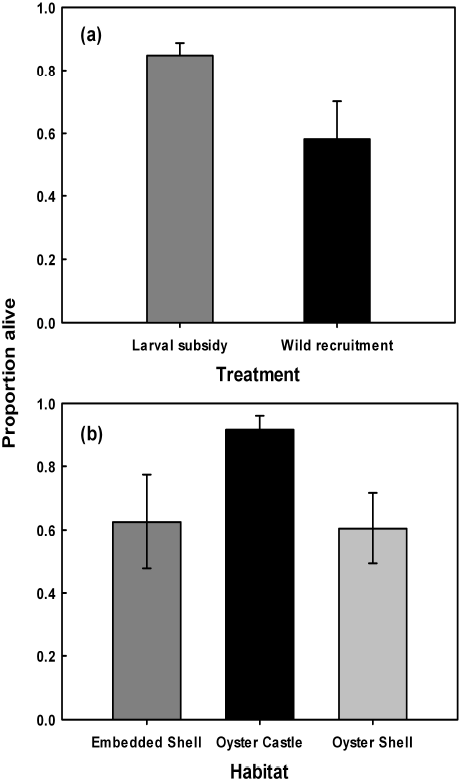
(a) Mean proportion of live recruits on experimental and control blocks, and (b) mean proportion of live recruits on the various substrates in the field experiment. Vertical bars represent one standard error of the mean.

### 3.3. Effect of recruit size on survival

In the log-linear analysis including experiment as a factor, there was a significant interaction between Experiment and Reef type (AIC reduced by 14.6 with 2 df), so the analyses were run separately for the field and mesocosm experiments.

#### 3.3.1. Mesocosm experiment

The global log-linear model was significant (likelihood ratio *χ*^2^ = 131.8, df = 18, P << 0.001). The final reduced model with stepwise addition of effects using AIC did not include any significant 2-way interactions, and only a significant main effect of size (likelihood ratio *χ*^2^ = 513.5, df = 20, P < 0.001). Overall, the probability of survival was 0.987 (Figure 5). The only substantial difference in survival was due to size, with juveniles greater than 30 mm shell length having much more than an order of magnitude greater probability of survival than juveniles less than 30 mm shell length (odds ratio = 24.7). Of the 25 dead juveniles, only 1 was greater than 30 mm.

#### 3.3.2. Field experiment

The global log-linear model was significant (likelihood ratio *χ*^2^ = 95.3, df = 18, P < 0.001). The final reduced model with stepwise addition of effects using AIC did not include any significant 2-way interactions, so the effect sizes were generated with the significant additive main effects model (likelihood ratio *χ*^2^= 34.1, df = 14, P << 0.005). Overall, the probability of mortality was 0.063 (Figure 8), which was nearly 5x higher than that in the mesocosms (0.013). The greatest difference in survival was due to size, with juveniles greater than 30 mm shell length having more than an order of magnitude greater probability of survival than juveniles less than 30 mm shell length (odds ratio = 12.6). The effect of reef type was also significant with oysters on OC having nearly fivefold higher survival probability than those on OS (odds ratio = 4.9) and ES (odds ratio = 4.4); there was no difference between ES and OS (odds ratio = 1.1). There was also a moderate effect of Subsidy increasing survival nearly threefold (odds ratio = 2.6).

**Figure 8:**
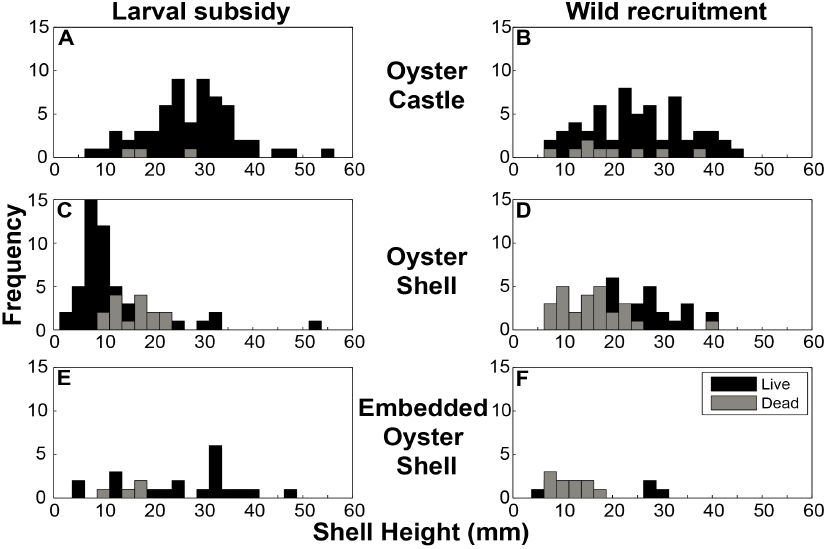
Size frequency histogram of live and dead oysters on Oyster Castles® (a & b), oyster shell (c & d), and embedded shell (e & f) in the field experiment.

## 4. Discussion

Although there have been numerous studies demonstrating that oysters can recruit to and survive on artificial oyster reefs (Soniat and Burton, 2005; Lipcius and Burke, 2006; Nestlerode et al., 2007; Burke, 2010; La Peyre et al., 2014; Dunn et al., 2014), this is the first to test oyster settlement and survival experimentally (i.e., both field and mesocosm experiments) on alternative reef substrates (i.e. Oyster Castles®), embedded oyster shell, and unconsolidated oyster shell reefs). In field and mesocosm experiments, alternative reef substrates recruited, retained, and hosted a greater oyster biomass than unconsolidated and embedded oyster shell. This study thus confirms that artificial reef materials are suitable alternative substrates for oyster restoration. Additionally, this study is among the first to demonstrate that field setting of hatchery-reared oyster larvae onto artificial substrates can be effective under controlled environmental conditions, thereby allowing for the possibility of enhancing settlement in locations that are limited by recruitment (Burke, 2010).

### 4.1. Alternative Substrates

Alternative reef substrates represent an effective means to enhance recruitment and survival of oysters in restoration efforts. In our mesocosm experiment, the Oyster Castles® recruited and hosted oyster density and biomass that were double that of unconsolidated shell; those on unconsolidated shell were nearly twice that of embedded shell. These differences were due to recruitment, rather than survival, because survival was high in all treatments and only differed by less than 3% across the three reef types. In the field experiment, the Oyster Castles® and unconsolidated shell were approximately equivalent in total recruit density, with the Oyster Castles® and unconsolidated shell retaining a total recruit density that was threefold higher than that on embedded shell. Live recruit density and biomass were about fivefold higher on Oyster Castles® than on embedded shell; on unconsolidated shell they were slightly greater than that on embedded shell. The reduction in live recruit density on unconsolidated shell was due to the low survival of recruits—60% on unconsolidated shell and 91% on Oyster Castles®.

The performance of the Oyster Castles® was likely due to the enhanced vertical relief of the substrate (14 cm on Oyster Castles® compared to 7 cm for embedded shell and 4 cm for unconsolidated shell), despite approximately equivalent surface area between reef treatment types. The higher relief would have allowed a greater percentage of substrate surface to remain above the sediment, out of hypoxic conditions, free from siltation, and in a fixed location. High levels of siltation were observed on the fixed embedded shell, whereas unconsolidated shells could be turned during periods of high wave activity (e.g., storm events), cleansed of silt, and remain exposed for settlement. Despite the propensity of unconsolidated shell to keep settlement surfaces exposed for recruitment, the turning of unconsolidated shells also repositioned some existing recruits into suboptimal orientations (i.e., inducing burial of recruits) yielding reduced survival of recruits (60% on unconsolidated shell). The enhanced vertical relief provided by Oyster Castles® allowed for settlement surfaces to remain exposed, normoxic, and fixed in location, which, in part, allowed for a fivefold higher oyster density and a fivefold higher probability of survival on Oyster Castles® relative to unconsolidated shell and embedded shell.

Elevation above the benthos, however, is not the only probable mechanism driving the differences in oyster density and biomass between reef treatment types. Unconsolidated shell provided both a greater amount of interstitial space for oyster settlement and a greater amount of rugose bottom area relative to the embedded shell treatments, which consisted of a horizontal base surface of concrete into which oyster shells were embedded. The combination of these two factors likely explains how the unconsolidated shell hosted an oyster density and biomass that was double that of embedded shell in the mesocosm experiment. Oyster Castles® provide interstitial space, a large amount of surface area for settlement, and heightened vertical relief, all of which combine to enhance oyster recruitment and biomass. Despite the relative failure of the remote setting in the field experiment to artificially enhance recruitment on the various reef substrates, the Castles® exhibited sufficient capacity to attract wild settlers at a level equivalent to, or greater than, unconsolidated shell.

### 4.2. Remote Setting

Field setting of larvae may be a useful approach to enhance bivalve recruitment in restoration efforts. In our mesocosm experiment, recruit density was about 50% higher in larval subsidy treatments, while biomass was more than twice as high. Recruits from the larval subsidy settled earlier and grew larger than wild recruits, which resulted in a greater fraction of juveniles greater than 30 mm shell length in larval subsidy treatments. Moreover, within the field experiment, the proportion of live recruits was higher among larval subsidy treatments, which increased survival by 47%. This increase in survival amongst larval subsidy treatments is likely due to the “swamping” of local predators by the pulse of larval subsidy recruits (Seitz et al., 2001). This partial prey refuge may allow a greater portion of these juveniles to escape post-settlement predation (Newell et al., 2007) and to grow to a size greater than 30 mm shell length where they have more than an order of magnitude greater probability of survival than juveniles less than 30 mm shell length. Thus, efficient pre-seeding may serve an important role in bolstering the resiliency of reef substrates by enhancing juvenile oyster survival (Burke, 2010).

Previous studies that have utilized field remote setting methods have yielded success in maximizing bivalve shellfish recruitment. Arnold et al. (2005) utilized a coupled mesocosm-field study design to examine the efficacy of remote setting of laboratory-reared bay scallops (*Argopecten irradians* Lamarck 1819) into controlled nurseries with subsequent transplantation into field locations and concluded that planting of cultured bay scallops was a successful strategy for increasing spawning stock density. Leverone et al. (2010) directly released hatchery-reared bay scallop larvae into two West Florida estuaries and concluded that larval remote setting can serve as an effective means to increase local scallop recruitment. In a 2006 VIMS study, a silt fence enclosure was used effectively to deter remotely set triploid oyster larvae from escaping the experimental replicates; only 1 in 60 oysters sampled outside the experimental replicates was a triploid (Burke, 2010). Thus, under favorable field conditions, remote setting of hatchery-reared oyster larvae onto reef substrates could serve as an effective restoration strategy.

The mesocosms afforded optimal hydrodynamic conditions allowing for concomitant increases in oyster recruitment, growth, and survival relative to the field experiment; there was 5x greater recruitment, 10x greater biomass, and nearly 5x greater survival (probability of mortality reduced from 0.063 to 0.013). Given that the greatest difference in oyster survival within the field experiment was due to size (i.e., juveniles greater than 30 mm shell length having more than an order of magnitude greater probability of survival than juveniles less than 30 mm shell length), remote setting of larvae onto reef substrates within mesocosms with growth to a mean juvenile size greater than 30 mm shell length and subsequent transplantation into field restoration locations could serve to dramatically enhance local spawning stock biomass, especially in areas that are recruitment-limited.

### 4.3. Implications for Oyster Restoration

Oyster Castles®, along with other similar alternative reef substrates continue to be utilized in both intertidal and subtidal reef restoration efforts by various governmental and nongovernmental agencies along the Atlantic and Gulf coasts, including a restoration effort conducted by the authors within a tributary of Chesapeake Bay (Figure 9) (McBride, 2012). These reef substrates are suitable for use by municipalities, commercial landowners, and individual homeowners interested in shoreline stabilization (i.e., to attenuate wave energy) and erosion control (i.e., accretion of shoreline behind reef structure), especially given their ability to be constructed from readily available, inexpensive materials. The performance measures for alternative reef materials reported in this study will be essential for comparison and assessment of current and future restoration projects, which should consider use of alternative reef substrates and field setting of larvae to maximize oyster recruitment while simultaneously minimizing the cost of reef restoration.

**Figure 9:**
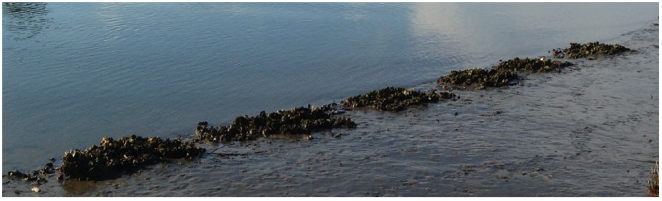
Photograph captured five years post-deployment of a reef constructed using substrates from the mesocosm and field experiments within the intertidal zone of Indian Field Creek, a tributary of Chesapeake Bay.

## 5. Acknowledgements

We thank the VIMS Marine Conservation Biology and Community Ecology Labs and VIMS 2008 Governor’s School interns for their assistance. We also thank The Nature Conservancy, Allied Concrete Corporation, and the VIMS Aquaculture Genetics and Breeding Technology Center (ABC) for providing necessary materials. This is Contribution XXXX from the Virginia Institute of Marine Science.

---

**Table S1:** Complete model selection results for the mesocosm experiment.

| Model | $AICc$ | $\Delta_i$ | $w_i$ | $r^2$ |
| --- | --- | --- | --- | --- |
| *(a) Total recruits* |  |  |  |  |
| $g_1$ | 172.8 | 10.9 | $< 0.01$ | 0.91 |
| $g_2$ | 169.2 | 7.2 | 0.02 | 0.90 |
| $g_3$ | 161.9 | 0.0 | 0.93 | 0.88 |
| $g_4$ | 168.3 | 6.3 | 0.04 | 0.78 |
| $g_5$ | 190.6 | 28.6 | $< 0.01$ | 0.09 |
| *(b) Live recruits* |  |  |  |  |
| $g_1$ | 172.6 | 11.0 | $< 0.01$ | 0.91 |
| $g_2$ | 168.9 | 7.3 | 0.02 | 0.89 |
| $g_3$ | 161.6 | 0.0 | 0.93 | 0.87 |
| $g_4$ | 167.9 | 6.2 | 0.04 | 0.78 |
| $g_5$ | 189.6 | 27.9 | $< 0.01$ | 0.10 |
| *(c) Live recruit biomass* |  |  |  |  |
| $g_1$ | 127.1 | 12.5 | $< 0.01$ | 0.73 |
| $g_2$ | 121.8 | 7.2 | 0.02 | 0.71 |
| $g_3$ | 114.6 | 0.0 | 0.90 | 0.66 |
| $g_4$ | 120.2 | 5.7 | 0.05 | 0.42 |
| $g_5$ | 121.6 | 7.1 | 0.03 | 0.24 |
| *(d) Proportion of live recruits* |  |  |  |  |
| $g_1$ | 86.0 | 16.4 | $< 0.01$ | 0.72 |
| $g_2$ | 90.1 | 12.4 | $< 0.01$ | 0.67 |
| $g_3$ | 99.6 | 2.9 | 0.19 | 0.66 |
| $g_4$ | 102.4 | 0.0 | 0.81 | 0.64 |
| $g_5$ | 87.7 | 14.7 | $< 0.01$ | 0.02 |

**Table S2:**
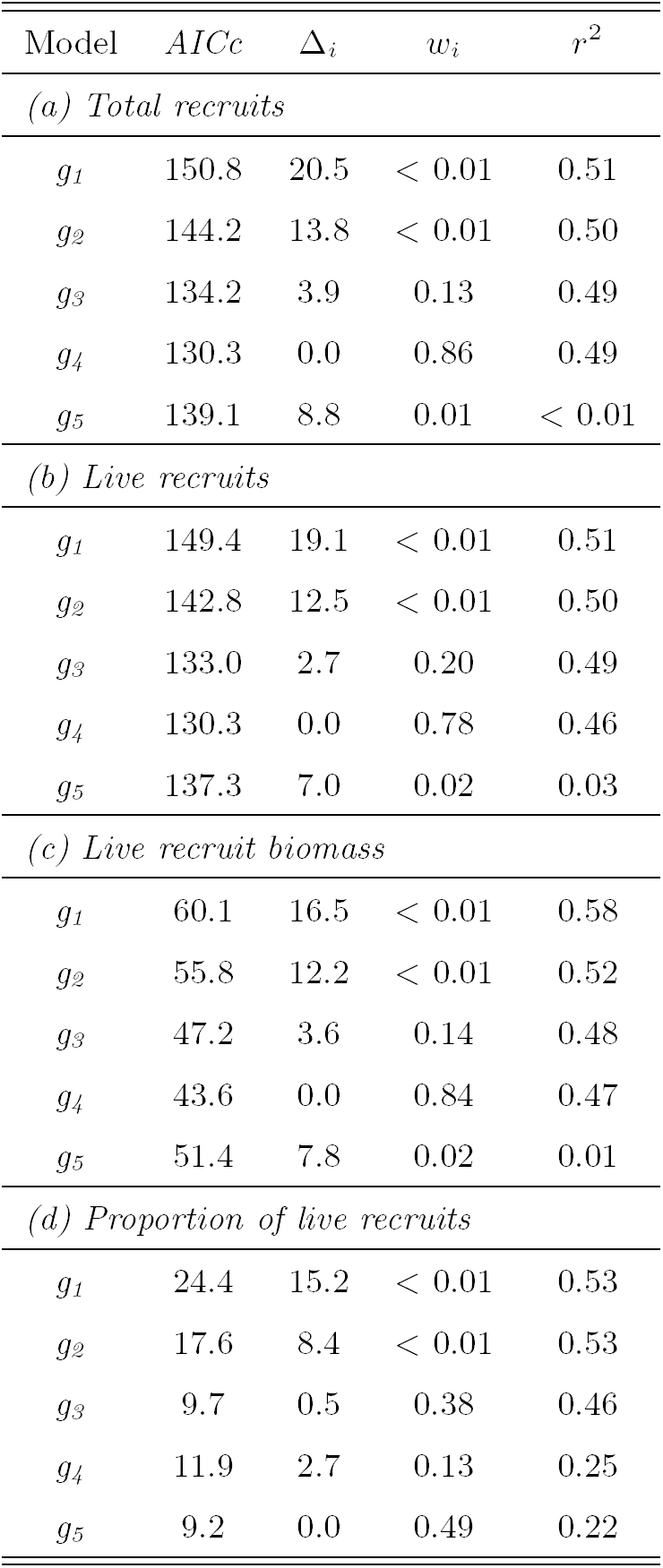
Complete model selection results for the field experiment.

| Model | $AICc$ | $\Delta_i$ | $w_i$ | $r^2$ |
| --- | --- | --- | --- | --- |
| *(a) Total recruits* |  |  |  |  |
| $g_1$ | 150.8 | 20.5 | $< 0.01$ | 0.51 |
| $g_2$ | 144.2 | 13.8 | $< 0.01$ | 0.50 |
| $g_3$ | 134.2 | 3.9 | 0.13 | 0.49 |
| $g_4$ | 130.3 | 0.0 | 0.86 | 0.49 |
| $g_5$ | 139.1 | 8.8 | 0.01 | $< 0.01$ |
| *(b) Live recruits* |  |  |  |  |
| $g_1$ | 149.4 | 19.1 | $< 0.01$ | 0.51 |
| $g_2$ | 142.8 | 12.5 | $< 0.01$ | 0.50 |
| $g_3$ | 133.0 | 2.7 | 0.20 | 0.49 |
| $g_4$ | 130.3 | 0.0 | 0.78 | 0.46 |
| $g_5$ | 137.3 | 7.0 | 0.02 | 0.03 |
| *(c) Live recruit biomass* |  |  |  |  |
| $g_1$ | 60.1 | 16.5 | $< 0.01$ | 0.58 |
| $g_2$ | 55.8 | 12.2 | $< 0.01$ | 0.52 |
| $g_3$ | 47.2 | 3.6 | 0.14 | 0.48 |
| $g_4$ | 43.6 | 0.0 | 0.84 | 0.47 |
| $g_5$ | 51.4 | 7.8 | 0.02 | 0.01 |
| *(d) Proportion of live recruits* |  |  |  |  |
| $g_1$ | 24.4 | 15.2 | $< 0.01$ | 0.53 |
| $g_2$ | 17.6 | 8.4 | $< 0.01$ | 0.53 |
| $g_3$ | 9.7 | 0.5 | 0.38 | 0.46 |
| $g_4$ | 11.9 | 2.7 | 0.13 | 0.25 |
| $g_5$ | 9.2 | 0.0 | 0.49 | 0.22 |

## References

Anderson, D. R. (2008). Model based inference in the life sciences: a primer on evidence. Springer.

Arnold, W. S., Blake, N. J., Harrison, M. M., Marelli, D. C., Parker, M. L., Peters, S. C., and Sweat, D. E. (2005). Restoration of bay scallop (*Argopecten irradians*(Lamarck)) populations in Florida coastal waters: planting techniques and the growth, mortality and reproductive development of planted scallops. Journal of Shellfish Research, 24(4):883–904.

Bonar, D., Coon, S., Walch, M., Weiner, R., and Fitt, W. (1990). Control of oyster settlement and metamorphosis by endogenous and exogenous chemical cues. Bulletin of Marine Science, 46(2):484–498.

Burke, R. (2010). Alternative substrates as a native oyster (Crassostrea virginica) reef restoration strategy in Chesapeake Bay. School of Marine Science, College of William and Mary Doctoral Dissertation. Doctoral Dissertation.

Congrove, M. S., Wesson, J. A., and Allen, S. K. (2008). A practical manual for remote setting in Virginia. Virginia Sea Grant.

Dunn, R. P., Eggleston, D. B., and Lindquist, N. (2014). Effects of substrate type on demographic rates of eastern oyster (Crassostrea virginica). Journal of Shellfish Research, 33(1):177–185.

Haywood III, E., Soniat, T., and Broadhurst III, R. (1999). Alternatives to clam and oyster shell as cultch for eastern oysters. In Oyster Reef Restoration: A Synopsis and Synthesis of Approaches, pages 295–304. Virginia Institute of Marine Science Press.

Kennedy, V., Breitburg, D., Christman, M., Luckenbach, M., Paynter, K., Kramer, J., Sellner, K., Dew-Baxter, J., Keller, C., and Mann, R. (2011). Lessons learned from efforts to restore oyster populations in Maryland and Virginia, 1990 to 2007. Journal of Shellfish Research, 30(3):719–731.

Kennedy, V., Newell, R., and Eble, A. (1996). The Eastern Oyster (Crassostrea virginica). Maryland Sea Grant.

La Peyre, M., Furlong, J., Brown, L. A., Piazza, B. P., and Brown, K. (2014). Oyster reef restoration in the northern Gulf of Mexico: Extent, methods and outcomes. Ocean & Coastal Management, 89:20–28.

Lenihan, H. S., Micheli, F., Shelton, S., and Peterson, C. H. (1999). The influence of multiple environmental stressors on susceptibility to parasites: an experimental determination with oysters. Limnology and Oceanography, 44:910–924.

Leverone, J. R., Geiger, S. P., Stephenson, S. P., and Arnold, W. S. (2010). Increase in bay scallop (*Argopecten irradians*) populations following releases of competent larvae in two west Florida estuaries. Journal of Shellfish Research, 29(2):395–406.

Lipcius, R. N. and Burke, R. P. (2006). Abundance, biomass and size structure of eastern oyster and hooked mussel on a modular artificial reef in the Rappahannock River, Chesapeake Bay. Special Report in Applied Marine Science and Ocean Engineering, 390.

McBride, S. (2012). Palmetto Castle, volume 4, pages 10–11. U.S.Army Corps of Engineers, Charleston District.

Nestlerode, J. A., Luckenbach, M. W., and O’Beirn, F. X. (2007). Settlement and survival of the oyster *Crassostrea virginica*on created oyster reef habitats in Chesapeake Bay. Restoration Ecology, 15(2):273–283.

Newell, R. I., Kennedy, V. S., and Shaw, K. S. (2007). Comparative vulnerability to predators, and induced defense responses, of eastern oysters *Crassostrea virginica* and non-native Crassostrea ariakensis oysters in Chesapeake Bay. Marine Biology, 152(2):449–460.

Newell, R. I. E. (1988). Ecological changes in Chesapeake Bay: are they the result of overharvesting the eastern oyster (*Crassostrea virginica*)? In Lynch, M. and Krome, E., editors, Understanding the Estuary, Advances in Chesapeake Bay Research, pages 536–546. Chesapeake Research Consortium, Gloucester Point, VA.

Osman, R. W. and Abbe, G. E. (1994). Post-settlement factors affecting oyster recruitment in the Chesapeake Bay, USA. In Dyer, K. R. and Orth, J., editors, Changes in Fluxes in Estuaries: Implications from Science to Management. Proceedings of Estuarine Research Federation Symposium, 13-18 September 1992, pages 335–340. Institute of Marine Studies, University of Plymouth.

Powers, S. P., Peterson, C. H., Grabowski, J. H., Lenihan, H. S., et al. (2009). Success of constructed oyster reefs in no-harvest sanctuaries: implications for restoration. Marine Ecology Progress Series, 389:159–170.

R Core Team (2014). R: A Language and Environment for Statistical Computing. R Foundation for Statistical Computing, Vienna, Austria.

Rothschild, B., Ault, J., Goulletquer, P., and Heral, M. (1994). Decline of the Chesapeake Bay oyster population: A century of habitat destruction and overfishing. Marine Ecology Progress Series, 111:29–39.

Ruesink, J., Lenihan, H., Trimble, A., Heiman, K., Micheli, F., Byers, J., and Kay, M. (2005). Introduction of non-native oysters: ecosystem effects and restoration implications. Annual Review of Ecology, Evolution, and Systematics, 36(1):643.

Schulte, D. M., Burke, R. P., and Lipcius, R. N. (2009). Unprecedented restoration of a native oyster metapopulation. Science, 325:1124–1128.

Seitz, R. D., Lipcius, R. N., Hines, A. H., and Eggleston, D. B. (2001). Density-dependent predation, habitat variation, and the persistence of marine bivalve prey. Ecology, 82(9):2435–2451.

Soniat, T. M., Broadhurst, R., and Haywood III, E. (1991). Alternatives to clam shell as cultch for oysters and the use of gypsum for the production of cultchless oysters. Journal of Shellfish Research, 10:405–410.

Soniat, T. M. and Burton, G. M. (2005). A comparison of the effectiveness of sandstone and limestone as cultch for oysters, Crassostrea virginica. Journal of Shellfish Research, 24(2):483–485.

Taylor, J. and Bushek, D. (2008). Intertidal oyster reefs can persist and function in a temperate North American Atlantic estuary. Marine Ecology Progress Series, 361:301.

Underwood, A. (1997). Experiments in Ecology: Their Logical Design and Interpretation Using Analysis of Variance. Cambridge University Press.

Wilberg, M., Livings, M., Barkman, J., Morris, B., and Robinson, J. (2011). Overfishing, disease, habitat loss, and potential extirpation of oysters in upper Chesapeake Bay. Marine Ecology Progress Series, 436:131–144.

